# Atlas for the Lateralized Visuospatial Attention Networks (ALANs): Insights from fMRI and Network Analyses

**DOI:** 10.1101/2024.02.13.580164

**Authors:** Loïc Labache, Laurent Petit, Marc Joliot, Laure Zago

## Abstract

Hemispheric specialization is central to human evolution and fundamental to human cognitive abilities. While being a defining feature of functional brain architecture, hemispheric specialization is overlooked to derive brain parcellations. Alongside language, which is typically lateralized in the left hemisphere, visuospatial attention is set to be its counterpart in the opposite hemisphere. However, it remains uncertain to what extent the anatomical and functional underpinnings of lateralized visuospatial attention mirror those supporting language. Building on our previous work, which established a lateralized brain atlas for language, we propose a comprehensive cerebral lateralized atlas delineating the anatomo-functional bases of visuospatial attention, ALANs. Combining task and resting-state functional connectivity analyses, we identified 95 lateralized brain areas comprising five networks supporting visuospatial attention processes. Among them, we can find two large-scale networks: the ParietoFrontal and TemporoFrontal networks. We identify hubs playing a pivotal role in the intra-hemispheric interaction within visuospatial attentional networks. The rightward lateralized ParietoFrontal encompasses one hub, the inferior frontal sulcus, while the TemporoFrontal network encompasses two right hubs: the inferior frontal cortex (pars triangularis and the anterior insula) and the posterior part of the superior temporal sulcus. Together, these networks encompass the homotope of the language network from the left hemisphere. This atlas of visuospatial attention provides valuable insights for future investigations into the variability of visuospatial attention and hemispheric specialization research. Additionally, it facilitates more effective comparisons among different studies, thereby enhancing the robustness and reliability of research in the field of attention.

---

**H**emispheric specialization is a fundamental principle in the functional organization of the human brain (Hervé et al., 2013). In about 90% of humans, who are righthanders, the left hemisphere is specialized for language and the motor control of their dominant hand (Labache et al., 2020, 2023; Mazoyer et al., 2014). In contrast, the right hemisphere is more dedicated to controlling visuospatial skills, including spatial attention (Hervé et al., 2013). This complementary hemispheric pattern between the language and spatial domain most likely results from evolutionary adaptive processes and selection pressure (Güntürkün & Ocklenburg, 2017; Heger et al., 2020). A significant contributor to this development and maintenance of hemispheric asymmetry is probably the corpus callosum, as suggested by Gazzaniga (Gazzaniga, 2000). However, the origin of the complementary patterns in hemispheric specialization is still a matter of debate (Francks, 2019; Gerrits, 2022; Thiebaut de Schotten et al., 2019; Tzourio-Mazoyer et al., 2020; Vingerhoets, 2019). Indeed, these complementary patterns remain misunderstood since they appear variable across the population, with a dependent relationship between language and spatial hemispheric lateralization only present in strongly left-handed individuals (Zago et al., 2016), while independence seems to be the rule for right-handed and mixed-handed individuals (Jia et al., 2021; Zago et al., 2016). This highlights the need to elaborate a normalized atlas to systematize the investigation of the lateralization of visuospatial processes at a regional level (Yeo & Eickhoff, 2016).

Although the identification of the neural attentional networks has been performed using various neuroimaging techniques in healthy individuals and patients with spatial neglect (Corbetta & Shulman, 2011; Petersen & Posner, 2012), the study of the lateralization has mainly been overlooked as compared to language (Hervé et al., 2013; Josse & Tzourio-Mazoyer, 2004; Mengotti et al., 2020; Tzourio et al., 1998). Visuospatial attention is a cognitive function traditionally lateralized to the right hemisphere (Heilman et al., 1993; Karnath & Rorden, 2012; Kinsbourne, 1970; Mesulam, 1999), as evidenced by the neuropsychological literature indicating spatial neglect after occipito-parietal lesions in the right hemisphere (Coppens et al., 2002; Dronkers & Knight, 1989; Suchan & Karnath, 2011). Unlike the lateralization of language, extensively studied and well-defined through established gold standard paradigms and techniques to explore its anatomo-functional bases, visuospatial functions lack a similar approach (Hervé et al., 2013). We showed that the line bisection judgment task is appropriate for investigating the anatomo-functional basis and the lateralization of brain regions involved in spatial attention in healthy participants (Zago et al., 2016, 2017).

A complex network of brain regions supports visuospatial attention. Neuropsychological studies differentiate attentional processes into two distinct types (Petersen & Posner, 2012): a slow, goal-oriented, and voluntary aspect, contrasted with a rapid, involuntary, stimulus-driven, and automatic element. The first one, the dorsal attentional network, encodes and sustains preparatory cues while modulating top-down sensory (visual, auditory, olfactory, and somatosensory) regions (Corbetta & Shulman, 2002). The second one, the ventral attentional network, activates when attention shifts to new, behaviorally significant events (Corbetta & Shulman, 2002). Key components of the dorsal network classically include the intraparietal sulcus, the superior parietal lobe, and the frontal eye fields at the junction between the superior frontal and precentral sulci. In contrast, the temporoparietal junction, the inferior part of the middle frontal gyrus, the inferior frontal gyrus, and the anterior insula constitute the core regions of the ventral network. In addition to these cortical structures, a set of subcortical structures, including the pulvinar, the superior colliculi, the head of caudate nuclei, and a group of brainstem nuclei, have been identified as involved in the organization of the ventral and dorsal attentional networks (Alves et al., 2022).

Despite the established roles of the dorsal and ventral attentional networks in visuospatial attention management, emerging discrepancies regarding their cerebral lateralization reveal a complex picture (Corbetta et al., 2000). Research indicates that visuospatial attention predominantly exhibits rightward lateralization during tasks (Petit et al., 2015; Schuster et al., 2017), yet the extent and direction of this lateralization remain subjects of debate. Notably, the dorsal attentional network is characterized by its bilateral operation in directing attention (Mengotti et al., 2020), with a slight leftward asymmetry at rest contrasted by a rightward asymmetry in its white matter pathways (Alves et al., 2022). Meanwhile, VAN’s bilateral rest activity further complicates our understanding of lateralization within these attentional frameworks (Alves et al., 2022; Mengotti et al., 2020). Finally, visuospatial attentional tasks also engaged executive and controlled processes subtended by prefrontal activations, rarely envisaged under the cerebral lateralization framework. While easily identified as distinct at rest (Gordon et al., 2016; Power et al., 2011; Yan et al., 2023), their naming and spatial topology are inconsistent across studies (Eickhoff et al., 2018; Uddin et al., 2023). Furthermore, task activation during attentional tasks does not respect the boundaries defined by rest, and part of each network can be seen activated conjointly (Corbetta & Shulman, 2011).

Here, leveraging multimodal approach (Hesling et al., 2019; Labache et al., 2019; Tzourio-Mazoyer et al., 2021), we aim to elucidate the anatomo-functional underpinnings and lateralization of visuospatial attentional networks. First, in a homogeneous sample of 130 right-handed individuals known for typical language lateralization, we identified significantly involved and lateralized brain regions in visuospatial attention processes using the line bisection judgment task. Second, we explored these identified brain areas’ network configuration and topological properties. This exploration is facilitated by applying agglomerative hierarchical clustering to resting-state data, enabling the extraction of distinct networks. Furthermore, we employed graph theory metrics to discern principal hubs integral to visuospatial attention processes. Finally, our study proposed an optimized model of visuospatial attention articulated through a lateralized atlas encompassing 95 well-characterized brain regions, the Atlas for Lateralized visuospatial Attentional Networks (ALANs). This model is a comprehensive framework for future research into the inter-individual variability of visuospatial attentional areas and the mechanisms underlying hemispheric specialization complementarity, enabling reproducible and reliable studies.

## Methods

### Participants

The study sample consisted of 130 participants from the BIL&GIN (Mazoyer et al., 2016) previously identified as typically brain-organized for language (Labache et al., 2020). The mean age of the sample was 27.3 years (*σ* = 6.3; range: 19–53 years; 64 women), and the mean level of education was 16.1 years (*σ* = 2.1 years; range: 11–20 years), corresponding to almost six years of education after the French baccalaureate. All participants were right-handed, as assessed with a mean Edinburgh score of +94.2 (*σ* = 10.3, (Oldfield, 1971)). All participants were free of brain abnormalities as assessed by a trained radiologist inspecting their structural T1-MRI scans. All participants gave their informed written consent and received compensation for their participation. The Basse-Normandie Ethics Committee approved the study protocol.

All participants completed a resting-state and two visuospatial task-related fMRI sessions, *i*.*e*., line bisection judgment and visually guided saccadic eye movements tasks. In the present study, we only report the line bisection judgment results.

### The Line Bisection Judgment Task

As detailed by Zago and colleagues (Zago et al., 2016), the line bisection judgment task consisted of a 2-s presentation of a horizontal line bisected by a short vertical line (subtending a visual angle of 1°), followed by a 10-s delay of a fixation cross. Participants decided whether the bisection mark was displayed precisely at the center of the horizontal line or slightly deviated to the left or the right. They responded with the right hand by pressing a three-button response pad with the index finger for answering “shifted to the left,” the middle finger for answering “centered,” and the ring finger for answering “deviated to the right.”

The horizontal line could be presented in three different positions along the horizontal axis (-7°, 0° or +7° of the center of the screen) and with three different lengths (6°, 7°, or 9° of visual angle). The bisection mark deviated by 0.3° on the center’s left or right of the center. Thirty-six trials were presented with an equal number of centered, leftward-, and rightward-bisected trials. A 12-s presentation of a fixation cross preceded and followed the first and last trials, respectively. Participants performed a practice phase before entering the scanner.

### Image Acquisition

Here, we report the main features of the structural and functional image acquisition previously described by Mazoyer and colleagues (Mazoyer et al., 2016).

#### Structural Image Acquisition

Images were acquired using a 3T Philips Intera Achieva scanner (Philips, Erlangen, The Netherlands). Structural imaging consisted of a high-resolution three-dimensional T1-weighted volume (T1w, sequence parameters: TR: 20 ms; TE: 4.6 ms; flip angle= 10°; inversion time: 800 ms; turbo field echo factor: 65; sense factor: 2; field of view: 256 x 256 x 180 mm^3^; 1 x 1 x 1 mm^3^ isotropic voxel size). The line between the anterior and posterior commissures was identified for each participant on a midsagittal section, and the T1MRI volume was acquired after orienting the brain in this bi-commissural coordinate system. T2*-weighted multislice images were also acquired (T2*-weighted fast field echo (T2*-FFE); sequence parameters: TR: 3.500 ms; TE: 35 ms; flip angle= 90°; sense factor: 2; 70 axial slices; 2 x 2 x 2 mm^3^ isotropic voxel size).

#### Functional Image Acquisition

Task-related functional volumes were acquired using a T2*-weighted echo-planar imaging sequence (T2*-EPI; TR: 2 s; TE: 35 ms; flip angle= 80°; 31 axial slices with a 240 x 240 mm^2^ field of view and 3.75 x 3.75 x 3.75 mm^3^ isotropic voxel size). The first four volumes of each sequence were discarded to allow for the stabilization of the MR signal.

Resting-state functional volumes were acquired as a single 8-minute-long run using the same T2*-EPI sequence (240 volumes) as the fMRI tasks. Before scanning, the participants were instructed to keep their eyes closed to relax, refrain from moving, stay awake, and let their thoughts come and go.

### Image Analysis

#### Functional Imaging Analysis for Task-Related and Resting-State Functional Volumes

As detailed by Labache and colleagues (Labache et al., 2019), for each participant, (1) the T2*-FFE volume was rigidly registered to the T1w; (2) the T1w volume was segmented into three brain tissue classes (grey matter, white matter, and cerebrospinal fluid); and (3) the T1w scans were normalized to the BIL&GIN template including 301 volunteers from the BIL&GIN database (aligned to the MNI space) using the SPM12 “normalize” procedure (http://www.fil.ion.ucl.ac.uk/spm/) with otherwise default parameters.

Data were corrected for slice timing differences for the three fMRI runs. The T2*-weighted volumes were realigned using a 6-parameter rigid-body registration to correct subject motion during the run. The participant T2*-EPI scans were then rigidly registered to the structural T2*-FFE image. Combining all registration matrices allowed warping the T2*-EPI functional scans from the subject acquisition space to the standard space (2 x 2 x 2 mm^3^ sampling size) with a single trilinear interpolation.

#### Specific Task-Related Functional Imaging Analysis

Statistical parametric mapping (SPM12, http://www.fil.ion.ucl.ac.uk/spm/) was used to process task-related fMRI data. First, a 6-mm full width at half maximum Gaussian filter was applied to volumes acquired during each run. Then, for each participant, the effects of interest were modeled by box-car functions computed with paradigm timing and convolved with a standard hemodynamic response function (SPM12). Individual contrast maps (line bisection judgment minus fixation) were calculated. These contrast maps (defined at the voxel level) were subjected to a region of interest analysis. BOLD signal variations were measured in 192 pairs of functionally defined regions of the AICHA atlas (Joliot et al., 2015) adapted to SPM12, excluding seven region pairs belonging to the orbital and inferior temporal parts of the brain in which signals were reduced due to susceptibility artifacts. For each participant, we computed this contrast map and calculated the right and left region BOLD signal variations for each of the 185 remaining pairs by averaging the contrast BOLD values of all voxels located within the region volume. The AICHA atlas was used here since it provides pairs of functionally homotopic regions and is thus well suited to measure functional asymmetries.

#### Specific Resting-State Functional Imaging Analysis

Time series of BOLD signal variations in white matter and cerebrospinal fluid (individual average time series of voxels that belonged to each tissue class) and temporal linear trends were removed from the rs-fMRI data series using regression analysis. Additionally, rs-fMRI data were bandpass filtered (0.01 Hz - 0.1 Hz) using a least-squares linear-phase finite impulse response filter design. For each participant and region, an individual BOLD rs-fMRI time series was computed by averaging the BOLD fMRI time series of all voxels within the region volume.

### Statistical Analysis

Statistical analysis was performed using R (R version: 4.2.2, (R Core Team, 2021)). Data wrangling was performed using the R library *dplyr* (R package version: 1.1.4, (Wickham et al., 2023)), and data visualization was performed using the R library *ggplot2* (R package version: 3.4.4, (Wickham, 2009)). Brain visualizations were realized using Surf Ice (NITRC: Surf Ice: Tool/resource Info, n.d.), and were made reproducible following guidelines to generate programmatic neuroimaging visualizations (Chopra et al., 2023).

We applied the three-step method previously developed by Labache and colleagues (Labache et al., 2019) to elaborate an atlas for the lateralized visuospatial attention networks. We will briefly outline this method in the subsequent sections.

#### Identification of the Anatomo-Functional Support of Visuospatial Attention

To identify the brain asymmetries underpinning the line bisection judgment task, we searched for regions that were significantly both activated and asymmetrical on average among the 130 participants for each hemisphere. We conducted a detailed conjunction analysis of the regions that exhibited significantly positive BOLD signal variations and higher values than their corresponding regions in the opposite hemisphere. A region was selected if it met two criteria: first, its mean *t*-value was positive, indicating significant activation in the right or left hemisphere at a significance threshold of *p* < 3.10^-4^, following the Bonferroni correction for multiple comparisons across 185 regions. Second, it demonstrated significant asymmetry at the same significance threshold. The overall significance threshold for these conjunction analyses was set at *p* = (3.10^-4^)^2^ = 7.10^-8^. This process was independently carried out for the left and right hemispheres, evaluating all 185 AICHA regions for both activation and asymmetry.

#### Network Organization of the Visuospatial Attention Regions

We first computed the intrinsic connectivity matrix for each participant (*n* = 130) to identify resting-state functional connectivity networks among the previously identified regions. The intrinsic connectivity matrix of off-diagonal elements were the Pearson correlation coefficients (*r*) between the rs-fMRI time series of region pairs (*n*_*region*_ = 95). The connectivity matrices were then Fisher *z*-transformed using the inverse hyperbolic tangent functions for each individual (R library psych; R package version: 2.3.9, (William Revelle, 2024)) before being averaged and *r*-transformed with the hyperbolic tangent function.

Second, based on the average connectivity matrix of the sample, we clustered the regions using an agglomerative hierarchical cluster analysis method (Sneath & Sokal, 1973; Ward, 1963). Each region was characterized according to its intrinsic connectivity pattern. Agglomerative hierarchical clustering was performed using Ward’s criterion as linkage criteria (Ward, 1963). Before classification, the average connectivity matrix was first transformed into a dissimilarity distance (*d*) using the following equation: *d* = (1 – *r*) / 2 (Doucet et al., 2011). The optimal number of clusters, determined using the R library *NbClust* (R package version: 1.1.4, (Charrad et al., 2014)), was found to be five. Based on 17 statistical indices, this method identified the most robust clustering scheme.

Finally, to evaluate the intrinsic inter-network communication, we computed the averaged temporal correlations between networks among the 130 participants. To determine the statistical significance of these correlations, we employed a non-parametric sign test with Bonferroni correction for multiple comparisons (10 comparisons), setting the adjusted significance level at *p* = 0.005.

#### Topological Characterization of Visuospatial Attention Networks

We applied graph theory to analyze intra-network communication across the five identified networks. Notably, we only included positive correlations in this analysis, as including negative correlations remains a debated topic in the field (Rubinov & Sporns, 2010).

We focused on two primary metrics to elucidate the network topology: degree and betweenness centrality. These metrics were instrumental in identifying hub regions, which are pivotal in influencing the overall network structure and flow of information.

Degree centrality was calculated for each region as the sum of its positive correlations with other regions within the same network. This measure effectively captures the overall connectedness of a region, highlighting its significance in the network. On the other hand, betweenness centrality quantifies the extent to which a region lies on the shortest paths between other regions. High betweenness centrality values indicate regions that act as essential bridges or intermediaries, facilitating communication across different network segments (Opsahl et al., 2010).

We adopted methodologies from Sporns and colleagues (Opsahl et al., 2010; Sporns et al., 2007) and van den Heuvel and colleagues (van den Heuvel et al., 2010) to determine hub regions. A region was classified as a hub if its degree and betweenness centrality values exceeded the mean and one standard deviation of these measures within the network’s regions (Labache et al., 2019). These identified hubs are crucial for maintaining network connectivity, enabling effective communication, and exerting substantial influence on the dynamics of information flow within the network.

## Results

### Identification of the Anatomo-Functional Support of Visuospatial Attention

We conducted a detailed conjunction analysis to identify the anatomical and functional bases of visuospatial attention. **Right hemisphere**. 66 regions met the selection criteria of being significantly activated in the right hemisphere and rightward asymmetrical (Figure 1). In the occipital lobe, rightward asymmetries were observed in various areas, including the calcarine (CAL3, CAL2), lingual (LING1, LING2, LING4, LING6), and fusiform parts (FUS4, FUS5, FUS6, FUS7), alongside the inferior (O3_2), middle (O2_1, O2_2, O2_3, O2_4), and lateral portions of the occipital gyri (Olat2, Olat4, Olat5), as well as the intraoccipital sulcus (ios).

**Figure 1.**
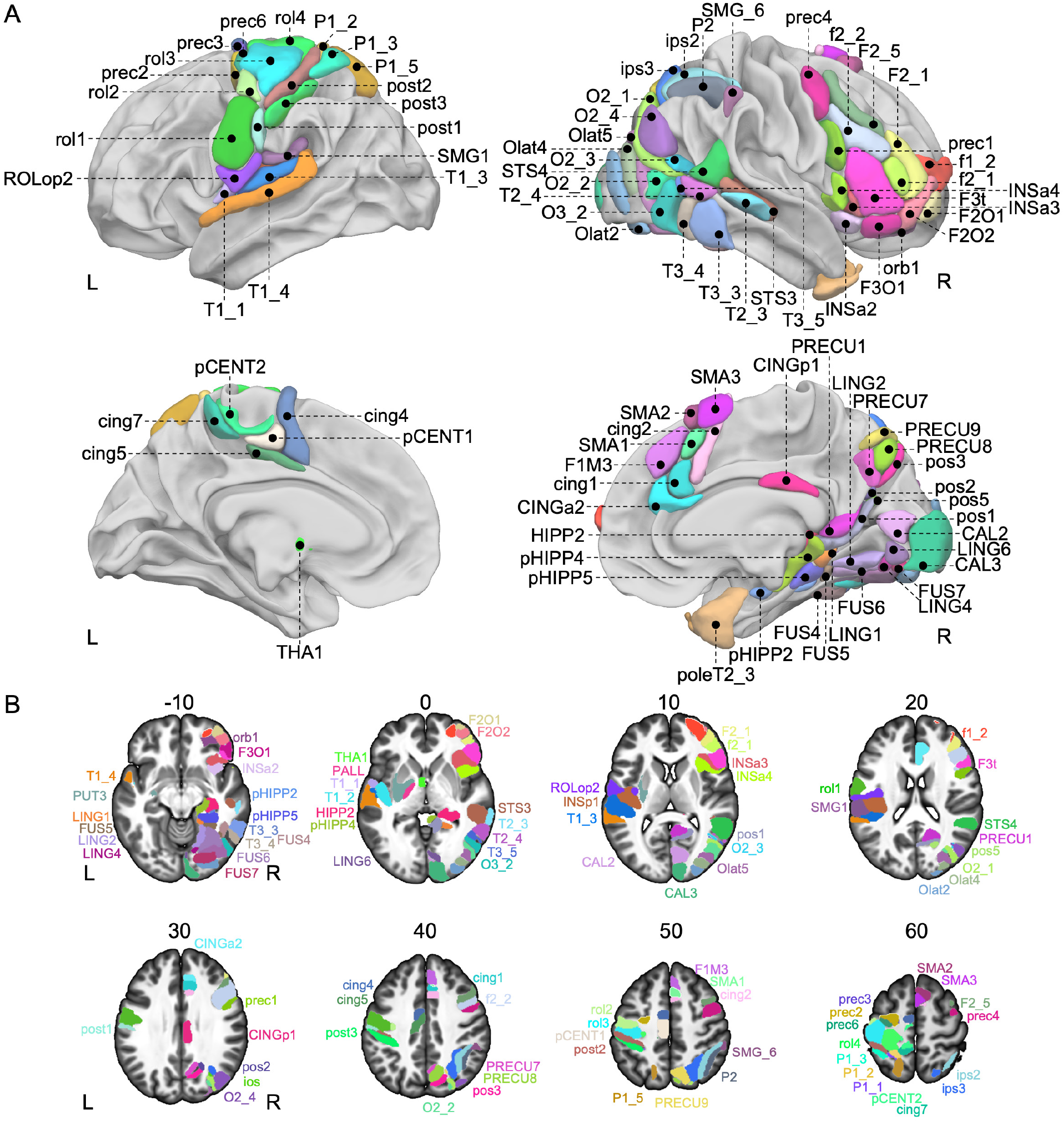
Asymmetric line bisection judgment-induced regions. **A**. View of the 66 right and 29 left AICHA regions on the 3D white surface rendering of the BIL&GIN display template (done in the MNI space with Surf Ice software (https://www.nitrc.org/projects/surfice/). Top row: Rightward line bisection judgment-regions showing conjoint BOLD activation in the right hemisphere and rightward asymmetry. Note that the intraoccipital sulcus (ios) is not visible in these views. Bottom row: Leftward line bisection judgment-regions showing conjoint BOLD activation in the left hemisphere and leftward asymmetry. Note that the posterior Insula (INSp1), the Putamen (PUT3), the Pallidum (PALL), the Superior Parietal (P1_1), the intraoccipital sulcus (ios), and the Superior Temporal Gyri (T1_2) are not visible in these views. **B**. Representation of these 95 regions on axial slices of the BIL&GIN display template with MRIcroGL software (https://www.nitrc.org/projects/mricrogl). The slices’ numbers correspond to the z-axis in the MNI space. Correspondences between the abbreviations and the full names of the AICHA atlas can be found in Table 1. Note that the right Temporal Pole (poleT2_3) is not visible on these axial slices. L: left; R: right. Abbreviations for the regions can be found in Table 1.

Within the parietal lobe, clusters of right-sided asymmetries were found in the intraparietal sulcus (ips2, ips3) and the inferior parietal gyrus (P2 and SMG6). On the medial surface, asymmetries were observed in different segments of the precuneus (PRECU1, PRECU7, PRECU8, PRECU9) along the parieto-occipital sulcus (pos1, pos2, pos3, pos5) extending towards the posterior part of the hippocampus (HIPP2) and parahippocampal formation (pHIPP2, pHIPP4, pHIPP5), as well as the anterior pole of the temporal gyrus (poleT2_3).

**Table 1.** Description of the 95 regions showing joint left activation and left asymmetry (resp. right activation and right asymmetry) during the Line Bisection Judgment task in 130 right-handers. The table displays the label of the network to which a region has been clustered, its abbreviation, its full anatomical name, the hemisphere to which it belongs, and the coordinates of its center of mass in MNI space. The number in parentheses corresponds to the functional subdivision of the region. The names of the regions correspond to the names defined in the AICHA atlas (Joliot et al., 2015).

| Network | Abbreviation | Region | Hemisphere | MNI coordinates |  |  |
| --- | --- | --- | --- | --- | --- | --- |
|  |  |  |  | X (mm) | Y (mm) | Z (mm) |
| VISU | CAL2 | Calcarine Gyrus (2) | Right | 10 | -78 | 9 |
|  | CAL3 | Calcarine Gyrus (3) | Right | 11 | -94 | 1 |
|  | FUS6 | Fusiform Gyrus (6) | Right | 29 | -62 | -9 |
|  | FUS7 | Fusiform Gyrus (7) | Right | 23 | -80 | -8 |
|  | LING2 | Lingual Gyrus (2) | Right | 21 | -60 | -6 |
|  | LING4 | Lingual Gyrus (4) | Right | 13 | -72 | -9 |
|  | LING6 | Lingual Gyrus (6) | Right | 7 | -79 | -3 |
|  | O2_2 | Middle Occipital Gyrus (2) | Right | 41 | -73 | 12 |
|  | O3_2 | Inferior Occipital Gyrus (2) | Right | 47 | -65 | -7 |
|  | Olat2 | lateral occipital Gyrus (2) | Right | 28 | -89 | -2 |
|  | Olat4 | lateral occipital Gyrus (4) | Right | 34 | -85 | 9 |
|  | Olat5 | lateral occipital Gyrus (5) | Right | 36 | -76 | 2 |
| SomatoMotor | cing4 | cingulate sulcus (4) | Left | -8 | -6 | 57 |
|  | cing5 | cingulate sulcus (5) | Left | -8 | -16 | 42 |
|  | cing7 | cingulate sulcus (7) | Left | -9 | -41 | 60 |
|  | INSp1 | Posterior Insula Gyrus (1) | Left | -42 | -19 | 14 |
|  | P1_1 | Superior Parietal Gyrus (1) | Left | -24 | -47 | 60 |
|  | P1_2 | Superior Parietal Gyrus (2) | Left | -19 | -47 | 68 |
|  | P1_3 | Superior Parietal Gyrus (3) | Left | -30 | -51 | 67 |
|  | P1_5 | Superior Parietal Gyrus (5) | Left | -16 | -61 | 61 |
|  | pCENT1 | Paracentral Lobule Gyrus (1) | Left | -7 | -17 | 51 |
|  | pCENT2 | Paracentral Lobule Gyrus (2) | Left | -10 | -29 | 66 |
|  | post1 | postcentral sulcus (1) | Left | -58 | -18 | 32 |
|  | post2 | postcentral sulcus (2) | Left | -41 | -33 | 55 |
|  | post3 | postcentral sulcus (3) | Left | -43 | -33 | 44 |
|  | prec2 | precentral sulcus (2) | Left | -25 | -8 | 59 |
|  | prec3 | precentral sulcus (3) | Left | -18 | -9 | 69 |
|  | prec6 | precentral sulcus (6) | Left | -30 | -11 | 65 |
|  | rol1 | Rolandic fissure (1) | Left | -54 | -8 | 32 |
|  | rol2 | Rolandic fissure (2) | Left | -44 | -14 | 51 |
|  | rol3 | Rolandic fissure (3) | Left | -39 | -23 | 61 |
|  | rol4 | Rolandic fissure (4) | Left | -23 | -29 | 65 |
|  | ROLOp2 | Rolandic Operculum (2) | Left | -51 | -9 | 14 |
|  | SMG1 | Supramarginal Gyrus (1) | Left | -54 | -30 | 21 |
|  | T1_1 | Superior Temporal Gyrus (1) | Left | -55 | -1 | 2 |
|  | T1_2 | Superior Temporal Gyrus (2) | Left | -45 | -11 | -2 |
|  | T1_3 | Superior Temporal Gyrus (3) | Left | -52 | -27 | 11 |
| PosteriorMedial | CINGp1 | Posterior Cingulate Gyrus (1) | Right | 5 | -26 | 29 |
|  | FUS4 | Fusiform Gyrus (4) | Right | 44 | -46 | -18 |
|  | FUS5 | Fusiform Gyrus (5) | Right | 32 | -47 | -11 |
|  | HIPP2 | Hippocampus Gyrus (2) | Right | 25 | -31 | -2 |
|  | ios | intraoccipital sulcus (1) | Right | 28 | -69 | 33 |
|  | ips3 | intraparietal sulcus (3) | Right | 26 | -62 | 46 |
|  | LING1 | Lingual Gyrus (1) | Right | 20 | -44 | -4 |
|  | O2_1 | Middle Occipital Gyrus (1) | Right | 36 | -74 | 25 |
|  | O2_4 | Middle Occipital Gyrus (4) | Right | 41 | -74 | 30 |
|  | pHIPP2 | Parahippocampal Gyrus (2) | Right | 29 | -25 | -19 |
|  | pHIPP4 | Parahippocampal Gyrus (4) | Right | 17 | -27 | -10 |
|  | pHIPP5 | Parahippocampal Gyrus (5) | Right | 27 | -36 | -12 |
|  | poleT2_3 | Middle Tempora Pole Gyrus (3) | Right | 26 | 6 | -36 |
|  | pos1 | parieto-occipital sulcus (1) | Right | 13 | -54 | 8 |
|  | pos2 | parieto-occipital sulcus (2) | Right | 16 | -61 | 26 |
|  | pos3 | parieto-occipital sulcus (3) | Right | 14 | -73 | 37 |
|  | pos5 | parieto-occipital sulcus (5) | Right | 21 | -66 | 20 |
|  | PRECU1 | Precuneus Gyrus (1) | Right | 13 | -53 | 14 |
|  | PRECU7 | Precuneus Gyrus (7) | Right | 7 | -63 | 36 |
|  | PRECU8 | Precuneus Gyrus (8) | Right | 11 | -68 | 41 |
|  | PRECU9 | Precuneus Gyrus (9) | Right | 13 | -68 | 50 |
|  | T3_3 | Inferior Temporal Gyrus (3) | Right | 57 | -46 | -14 |
|  | T3_4 | Inferior Temporal Gyrus (4) | Right | 54 | -58 | -11 |

**Table 1 |** (continued)
| Network | Abbreviation | Region | Hemisphere | MNI coordinates |  |  |
| --- | --- | --- | --- | --- | --- | --- |
|  |  |  |  | X (mm) | Y (mm) | Z (mm) |
| TemporoFrontal | cing1 | cingulate sulcus (1) | Right | 7 | 27 | 31 |
|  | cing2 | cingulate sulcus (2) | Right | 8 | 13 | 47 |
|  | CINGa2 | Anterior Cingulate Gyrus (2) | Right | 7 | 33 | 23 |
|  | F3O1 | Inferior Frontal Gyrus: Pars Orbitalis (1) | Right | 44 | 33 | -14 |
|  | F3t | Inferior Frontal Gyrus: Pars Triangularis (1) | Right | 50 | 29 | 5 |
|  | INSa2 | Anterior Insula Gyrus (2) | Right | 35 | 18 | -13 |
|  | INSa3 | Anterior Insula Gyrus (3) | Right | 37 | 24 | -0 |
|  | INSa4 | Anterior Insula Gyrus (4) | Right | 41 | 15 | 4 |
|  | O2_3 | Middle Occipital Gyrus (3) | Right | 45 | -63 | 15 |
|  | PALL | Pallidum (1) | Left | -19 | -8 | -1 |
|  | PUT3 | Putamen (3) | Left | -28 | -6 | 2 |
|  | SMA2 | Supplementary Motor Area (2) | Right | 11 | 18 | 63 |
|  | SMA3 | Supplementary Motor Area (3) | Right | 6 | 10 | 66 |
|  | STS3 | superior temporal sulcus (3) | Right | 53 | -32 | -0 |
|  | STS4 | superior temporal sulcus (4) | Right | 55 | -46 | 15 |
|  | T1_4 | Superior Temporal Gyrus (4) | Left | -59 | -23 | 4 |
|  | T2_3 | Middle Temporal Gyrus (3) | Right | 62 | -31 | -5 |
|  | T2_4 | Middle Temporal Gyrus (4) | Right | 57 | -53 | 3 |
|  | T3_5 | Inferior Temporal Gyrus (5) | Right | 49 | -58 | 4 |
|  | THA1 | Thalamus (1) | Left | -4 | 0 | 1 |
| ParietoFrontal | f1_2 | superior frontal sulcus (2) | Right | 28 | 56 | 7 |
|  | f2_1 | inferior frontal sulcus (1) | Right | 46 | 40 | 10 |
|  | f2_2 | inferior frontal sulcus (2) | Right | 44 | 19 | 28 |
|  | F1M3 | Medial Superior Frontal Gyrus (3) | Right | 6 | 33 | 45 |
|  | F2_1 | Middle Frontal Gyrus Gyrus (1) | Right | 41 | 44 | 13 |
|  | F2_5 | Middle Frontal Gyrus Gyrus (5) | Right | 42 | 17 | 41 |
|  | F2O1 | Middle Frontal Gyrus: Pars Orbitalis (1) | Right | 36 | 57 | -6 |
|  | F2O2 | Middle Frontal Gyrus: Pars Orbitalis (2) | Right | 40 | 50 | -4 |
|  | ips2 | intraparietal sulcus (2) | Right | 37 | -52 | 48 |
|  | orb1 | orbital sulcus (1) | Right | 26 | 41 | -15 |
|  | P2 | Inferior Parietal Gyrus (1) | Right | 43 | -53 | 48 |
|  | prec1 | precentral sulcus (1) | Right | 50 | 10 | 24 |
|  | prec4 | precentral sulcus (4) | Right | 44 | 1 | 48 |
|  | SMA1 | Supplementary Motor Area Gyrus (1) | Right | 6 | 21 | 49 |
|  | SMG6 | Supramarginal Gyrus (6) | Right | 54 | -38 | 44 |

In the temporal lobe, conjunction of activations and asymmetries were present in the lateral portions of the inferior (T3_3, T3_4, T3_5) and middle (T2_3, T2_4) temporal gyri, as well as in the superior temporal sulcus (STS4 and STS3). Moving to the frontal lobe, regions were identified in the inferior and orbital regions (F3O1, F3t, F2O1, F2O2, orb1), extending into the anterior insula (INSa2, INSa3, INSa4). Furthermore, right-brain-dominant asymmetries were observed in the precentral sulcus (prec4 and prec1) and various segments of the middle frontal gyrus (F2_1, F2_5), as well as the inferior and superior frontal sulci (f2_1, f2_2, f1_2). On the medial surface, asymmetries were detected in the supplementary motor area (SMA1, SMA2, SMA3), the median superior frontal gyrus (F1M3), and the anterior and posterior parts of the cingulate gyrus (CINGa2, CINGp1, cing1, cing2).

### Left hemisphere

A total of 29 regions met the selection criteria in this study (Figure 1). Notable asymmetries were observed on the lateral surface, specifically along the Rolandic sulcus (rol1, rol2, rol3, rol4), extending to the precentral sulcus (prec2, prec3, prec6), and the postcentral sulcus (post1, post2, post3) corresponding to the sensorimotor cortex. Leftward asymmetries were also observed in the Rolandic operculum (ROLop2), posterior insula (INSp1), and the lower part of the supramarginal gyrus (SMG1). Additionally, subcortical asymmetries favoring the left side were found in the pallidum (PALL), thalamus (THA1), and putamen (PUT3). The superior temporal gyrus (T1_1, T1_2, T1_3, T1_4) and superior parietal gyrus (P1_1, P1_2, P1_3, P1_5) exhibited leftward asymmetry. On the medial face, asymmetry was observed in the posterior sections of the cingulum (cing4, cing5, cing7), as well as two regions in the paracentral lobule (pCENT1, pCENT2).

Correspondences between the abbreviations and the full names of the regions can be found in Table 1.

### Network Organization of the Visuospatial

#### Attention Regions

To identify the network organization of visuospatial attention regions, we conducted agglomerative hierarchical clustering on the 95 regions previously identified through conjunction analysis. We then assessed the inter-network communication by examining the temporal correlation across these networks. The significance of these correlations was tested using a non-parametric sign test.

##### Description of the Visuospatial Attention Intrinsic Networks

The agglomerative hierarchical clustering analysis revealed five networks from the selected set of 95 asymmetric line bisection judgment-induced regions (Figure 2, Table 1).

**Figure 2.**
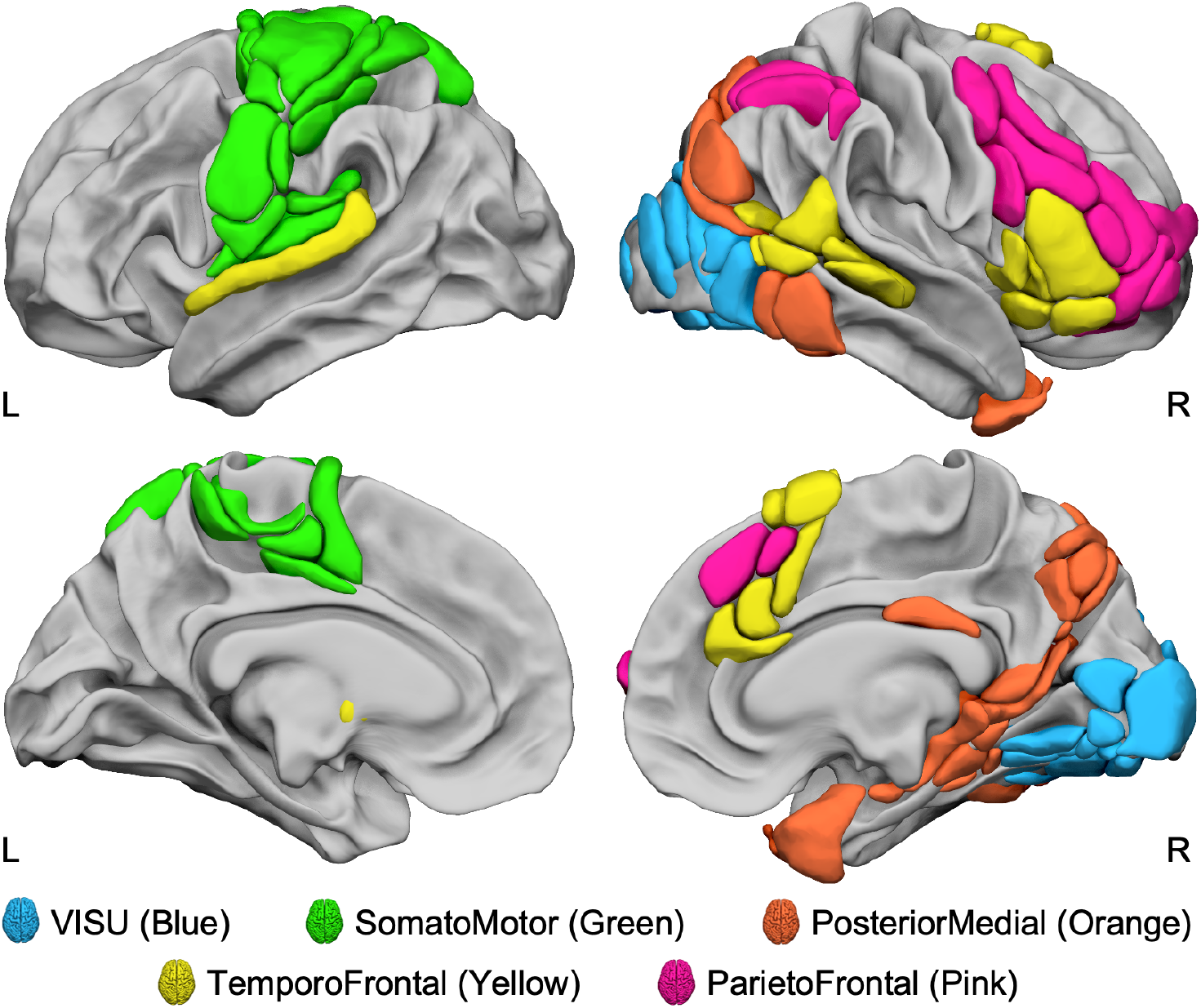
Lateral and medial views of the five intrinsic identified networks of the 95 regions asymmetrically involved in the line bisection judgment task, evidenced by the agglomerative hierarchical cluster analysis method. View of 3D white surfaces rendering on the BIL&GIN display template in the MNI space. L: left; R: right.

##### VISU network

This network includes 12 regions (Figure 2, in blue), all located bilaterally in the posterior part of the occipital lobe. We labeled it VISU because it aggregated regions acknowledged as involved in visual processing.

##### SomatoMotor network

This network includes most of the cortical regions found in the left hemisphere (Figure 2, in green). We labeled it SomatoMotor because it aggregated brain regions involved in the motor and somatosensory aspects of the response production (Tzourio-Mazoyer et al., 2021).

##### PosteriorMedial network

This third network encompasses 23 regions (Figure 2, in orange) located first on the medial surface, namely the dorsal medial parietal regions (precuneus and parieto-occipital sulcus) and the medial temporal regions (posterior part of the hippocampus and the parahippocampus), extending to the anterior fusiform and anterior temporal pole. Secondly, on the lateral surface, it aggregates the posterior part of the intraparietal (ips3) and intra-occipital sulci (ios), extending to the middle occipital gyrus (O2_1, O2_4) to the pole of the middle temporal (poleT2_3) and inferior temporal (T3_3, T3_4) gyri.

##### TemporoFrontal network

This network included 20 regions (Figure 2, in yellow), 16 being right-lateralized. On the right side, the temporoFrontal network aggregates all the regions located in the inferior and ventral frontal cortex (F3t, F3O1, INSa2, INSa3, INSa4) and the posterior part of the temporal cortex (STS4, STS3, T2_3, T2_4, T3_5) extending to the middle occipital gyrus (O2_3). On the medial wall, this network gathers most of the regions found in the supplementary motor area (SMA2, SMA3) and the anterior cingulate gyrus (cing1, cing2, CINGa2). On the left side, it aggregates the three subcortical regions (PALL, PUT3, and THA1) and the left superior temporal gyrus (T1_4).

##### ParietoFrontal network

This network consists of 15 rightward regions (Figure 2, in pink), predominantly located in the dorsal and anterior parts of the lateral frontal lobe. These regions include areas along the precentral sulcus (prec1, prec4) and the medial superior frontal cortex (SMA1, F1M3). The network also encompasses the inferior parietal lobe (SMG6, P2) and the intraparietal sulcus (ips_2).

##### Temporal Correlation Across Networks

The mean intrinsic connectivity analyses revealed the following correlations between the networks:

##### SomatoMotor and VISU networks

There was a positive correlation (*r* = 0.37) between the SomatoMotor and VISU networks, indicating that these two networks exhibit synchronized activity. The correlation was statistically significant (*p* = 1.10^-27^), suggesting a robust association between the two networks.

##### ParietoFrontal and TemporoFrontal networks

The ParietoFrontal and TemporoFrontal networks also showed a positive correlation (*r* = 0.29), indicating synchronized activity between these networks. The correlation was statistically significant (*p* = 7.10^-19^) suggesting a meaningful relationship between the ParietoFrontal and TemporoFrontal networks.

##### TemporoFrontal and ParietoFrontal with SomatoMotor and VISU networks

The TemporoFrontal network exhibited positive correlations with both the SomatoMotor network (*r* = 0.25, *p* = 1.10^-20^) and the VISU network (*r* = 0.14, *p* = 3.10^-10^), while the ParietoFrontal network showed negative correlations with them (SomatoMotor network, *r* = -0.11, *p* = 2.10^-5^; VISU network, *r* = -0.18, *p* = 3.10^-17^). It reveals a synchronized activity of the TemporoFrontal network and an antagonistic activity of the ParietoFrontal networks with the SomatoMotor and VISU networks.

##### PosteriorMedial with VISU and TemporoFrontal networks

The PosteriorMedial network positively correlated with the VISU network (*r* = 0.11, *p* = 3.10^-4^), indicating synchronized activity between these networks. However, there was a negative correlation between the PosteriorMedial and TemporoFrontal networks (*r* = -0.07, *p* = 5.10^-4^), suggesting potentially different functional characteristics or opposing activity patterns between these networks.

Notably, no significant correlation was found between the PosteriorMedial network and the SomatoMotor or ParietoFrontal networks (*p* > 0.40 for both), indicating a lack of strong associations between these specific network pairs.

### Topological Characterization of Visuospatial Attention Networks

We computed the degree and betweenness centrality to explore the topological organization of visuospatial attention networks and identify crucial hub regions.

The Degree Centrality (DC) values across the regions ranged from 1.36 to 6.18, while the standard deviation of DC remained consistently between 0.85 and 1.61. Similarly, the Betweenness Centrality (BC) values spanned from 0.32 to 4.17, displaying a range comparable to DC. Within the **ParietoFrontal network**, the hub significance thresholds were determined as 6.02 for DC (mean + *σ*) and 0.97 for BC (mean + *σ*). Only the inferior frontal sulcus region (f2_2, Figure 3) met the hub criteria, with a BC value of 1.35 (CI_95%_ = [1.06, 1.64]) and a DC value of 5.99 (CI_95%_ = [5.75, 6.22]).

**Figure 3.**
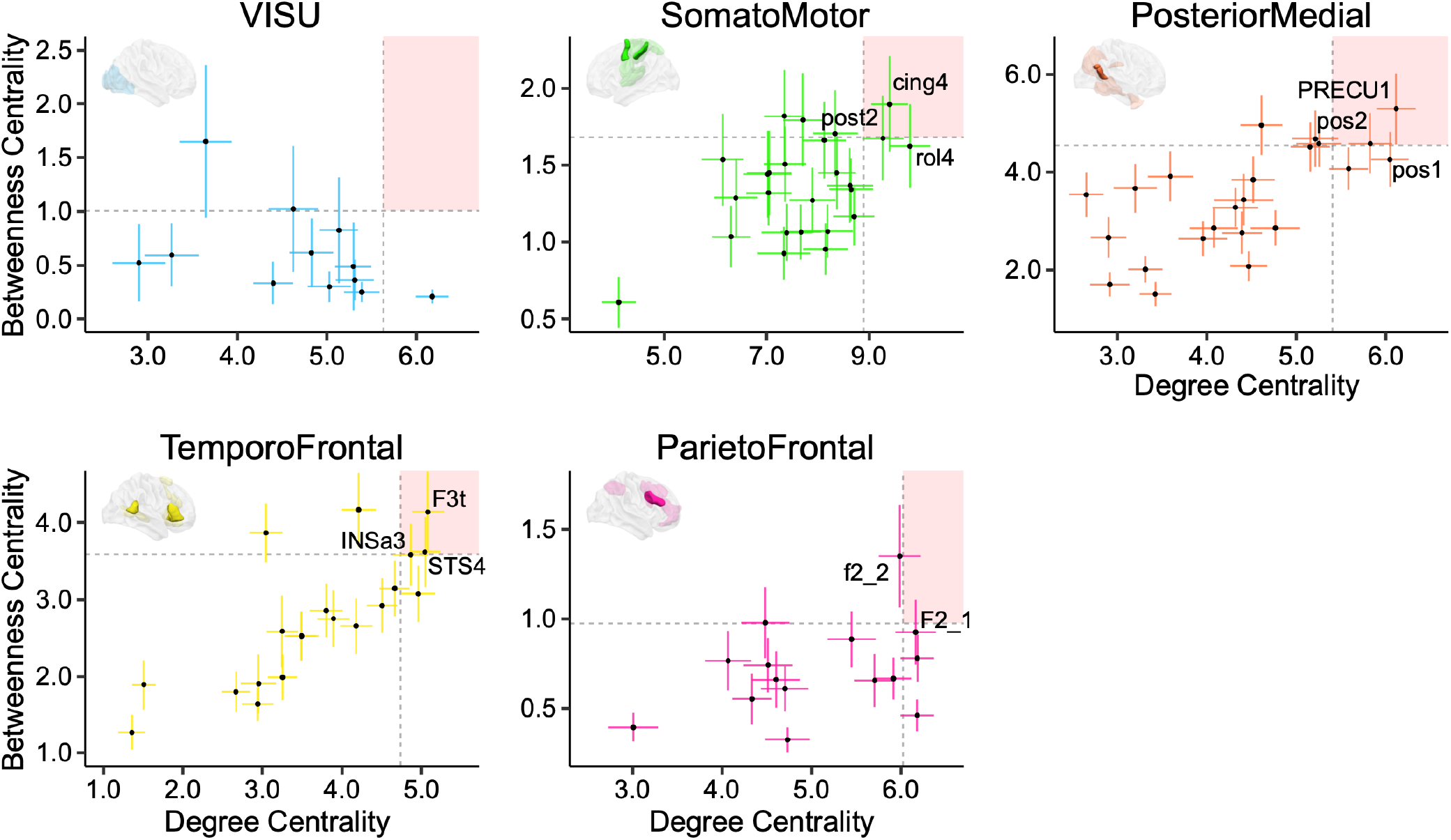
Identification of hubs. Plots of Degree Centrality (DC) versus Betweenness Centrality (BC) in each of the 5 networks. Bars are 95% confidence intervals for each DC and BC value of each region. The mean plus one standard deviation (*σ*) value of DC and BC defines the quadrants. regions located in the right superior quadrant are hubs and are illustrated on the corresponding hemisphere (solid regions). Abbreviations for the regions can be found in Table 1.

In the **TemporoFrontal network**, hubs were defined by thresholds of 3.59 for DC and 4.74 for BC. Three regions satisfied the hub definition: F3t (DC = 4.14, CI_95%_ = [3.09, 5.19]; BC = 5.08, CI_95%_ = [4.67, 5.49]), STS4 (DC = 3.62, CI_95%_ = [2.07, 4.54]; BC = 5.04, CI_95%_ = [4.65, 5.43]), and INSa3 (DC = 3.59, CI_95%_ = [2.78, 4.40]; BC = 4.86, CI_95%_ = [4.41, 5.31]) (Figure 3). INSa3 approached the BC hub threshold but still qualified as a hub.

In the **PosteriorMedial network**, PRECU1 (DC = 6.11, CI95% = [5.90, 6.33]; BC = 5.30, CI95% = [4.58, 6.02]) and pos2 (DC = 5.82, CI95% = [5.58, 6.07]; BC = 4.59, CI95% = [3.97, 5.20]) were identified as hubs, as their DC and BC values surpassed the set thresholds (DC ≥ 5.40, BC ≥ 4.55).

Within the **SomatoMotor network**, two sensorimotor regions were classified as hubs based on the thresholds of DC ≥ 8.90 and BC ≥ 1.68: post2 (DC = 9.27, CI95% = [8.86, 9.68]; BC = 1.68, CI95% = [1.40, 1.95]) and the neighboring cing4 in the medial wall (DC = 9.40, CI95% = [9.04, 9.76]; BC = 1.90, CI95% = [1.59, 2.21]).

None of the 12 regions in the **VISU network** met the chosen significance thresholds (DC ≥ 5.63, BC ≥ 1.00), thus not qualifying as hubs.

## Summary of the Results

In this study, we analyzed the activation and asymmetry of the brain in 130 right-handed participants engaged in a visuospatial attentional line bisection judgment task. Using the AICHA atlas, we identified 95 lateralized regions – 66 on the right and 29 on the left. Agglomerative hierarchical clustering based on intrinsic connectivity among these regions yielded five distinct intrinsic networks. These networks were named according to their anatomical locations: VISU, SomatoMotor, PosteriorMedial, TemporoFrontal, and ParietoFrontal.

Further analysis revealed notable intrinsic connectivity patterns. Strong positive correlations were observed between the SomatoMotor and VISU networks and between the ParietoFrontal and TemporoFrontal networks. The TemporoFrontal network also showed positive correlations with both the SomatoMotor and VISU networks. Conversely, the ParietoFrontal network exhibited negative correlations with the SomatoMotor and VISU networks.

Additionally, the PosteriorMedial network demonstrated positive correlations with the VISU network.

Graph metric analysis highlighted key hubs within these networks. Within the TemporoFrontal network, the right F3t, right INSa3, and right STS4 regions showed high degrees of centrality, indicating their significant roles as network hubs. The right-lateralized ParietoFrontal network’s lateral inferior frontal sulcus region (f2_2) also emerged as a prominent hub. In the PosteriorMedial network, the PRECU1 and pos2 regions located in the medial wall in the right precuneus and parieto-occipital sulcus were identified as hubs. Similarly, the pre-supplementary motor area (cing4) and the somatosensory cortex (post2) regions in the left-lateralized SomatoMotor network were also recognized as hubs. These hubs are pivotal in facilitating communication and the flow of information within their respective networks.

## Discussion

Our study identifies lateralized brain networks in a visuospatial attention task among a substantial sample of right-handed individuals, showing typical language organization. Using a multimodal approach, integrating the line bisection judgment task and resting-state acquisition, we identified 95 lateralized regions organized in five networks. Among these, two key rightward networks – the ParietoFrontal and TemporoFrontal – demonstrate strong synchronous fMRI signal oscillations at rest, organized around four core regions: the inferior frontal sulcus, the inferior frontal gyrus (pars triangularis), the anterior insula, and the posterior part of the superior temporal sulcus. Together, this work advances our understanding of organizing the anatomo-functional bases of visuospatial attention. It will also enable investigations into brain organization in atypical individuals and assess hemispheric complementarity mechanisms (Johnstone et al., 2020; Tzourio-Mazoyer et al., 2019; Vingerhoets, 2019).

In addition to the typical rightward functional asymmetries in temporoparietal and frontal regions, known to be recruited during visuospatial attentional task-related fMRI studies (Ciçek et al., 2009; Zago et al., 2016, 2017), we observed leftward asymmetries in relation to the somatomotor response production. These findings align with the typical brain functional organization, where visuospatial attention exhibits right-hemisphere dominance, and response production demonstrates left-hemisphere dominance in right-handers.

Among these lateralized brain regions recruited during the line bisection judgment task, the intrinsic connectivity analysis revealed the presence of five networks exhibiting spontaneous synchronization of low-frequency fMRI oscillations during rest. This analysis distinguished between local networks that clustered visual and somatomotor regions (VISU and SomatoMotor) and large-scale networks that clustered temporo-frontal regions (TemporoFrontal network), parieto-frontal regions (ParietoFrontal network), and posterior medial regions (PosteriorMedial network). This division aligns with other studies examining global brain intrinsic connectivity (Yeo et al., 2011; Gordon et al., 2016; Doucet et al., 2011).

To better characterize our LBJ-related lateralized networks, and as suggested by Uddin and colleagues (Uddin et al., 2023), we compared our five-network clustering to the seven-network parcellation proposed by Yan and colleagues (Yan et al., 2023), using a similar approach to Labache and colleagues (Labache et al., 2023). For each region of the AICHA atlas, we computed a distribution of overlap percentage with all seven canonical networks. Each region was assigned to the network with the greatest overlap (Figure 4.A). As depicted in Figure 4 (B and C), our five-network clustering approach revealed that the local visual and sensorimotor networks are concordant with those identified by Yan and colleagues, as evidenced by the significant overlap observed. For example, all regions clustered in the VISU network correspond to the Visual canonical network (Figure 4, in violet), and 72% of the regions in our SomatoMotor network align with the canonical Som/Motor network (Figure 4, in blue). Therefore, the overlap between our clustering approach and Yan’s parcellation is consistent for the local networks. The ParietoFrontal network overlaps with the Control network by 80% and the DorsAttn network by 20%, indicating that the rightward ParietoFrontal network groups together brain regions, subtending controlled and goal-oriented attentional processes. The overlap for the two other large-scale TemporoFrontal and PosteriorMedial networks is more scattered, which is consistent with the ongoing challenge in the existing literature to establish a consensus regarding the classification of different large-scale networks across various studies (Uddin et al., 2019; Witt et al., 2021). Each of these networks will be discussed in detail below.

**Figure 4.**
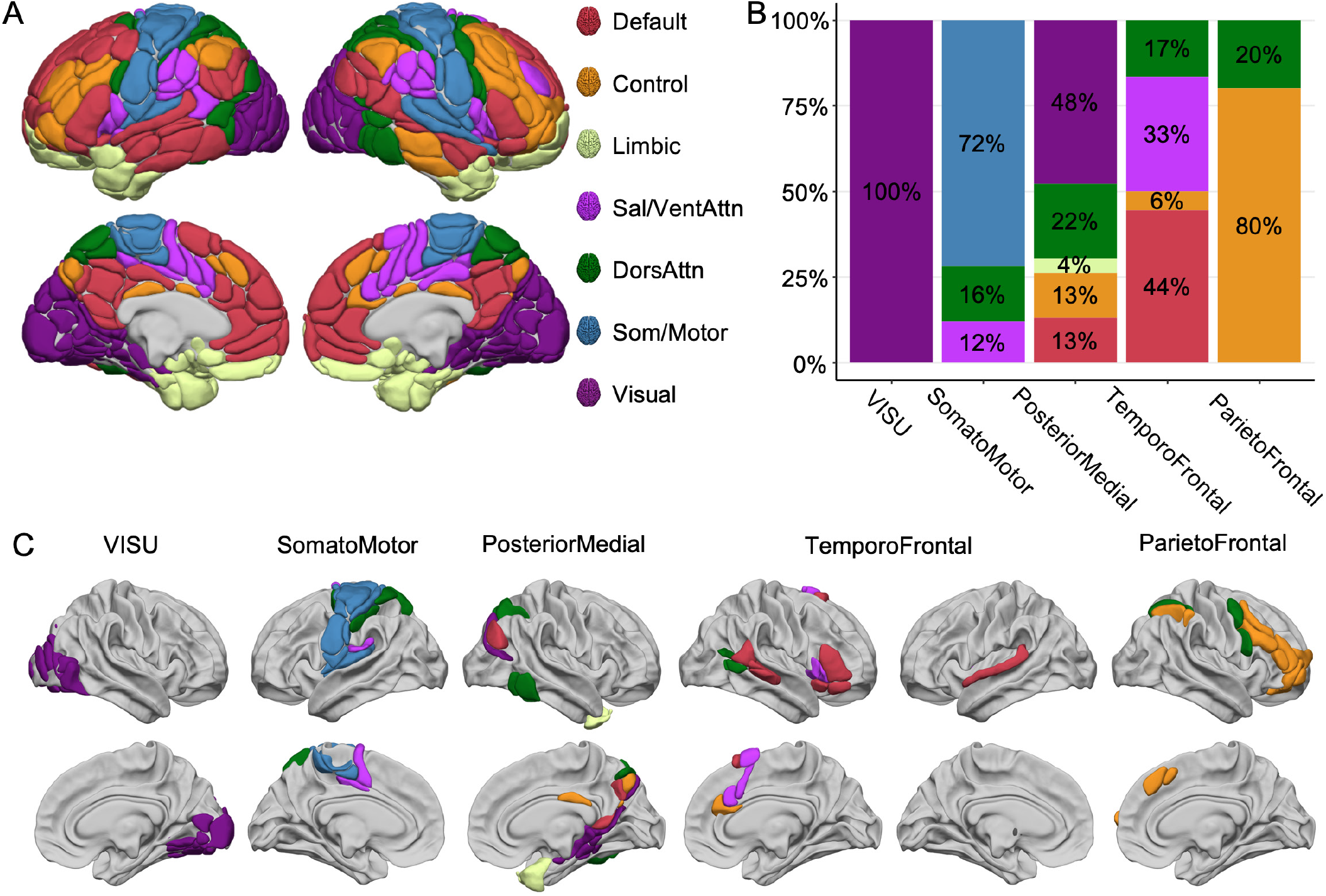
Comparison between the five ALANs (Atlas of Lateralized visuospatial Attentional Networks) clustered networks and the seven canonical network parcellation by Yeo and colleagues (Yeo et al., 2011) as proposed by Yan and colleagues (Yan et al., 2023). **A**. The 7-network parcellation is rendered on the AICHA atlas (Joliot et al., 2015). **B**. Repartition of the regions of the ALANs five-network parcellation across the seven canonical networks. Color code corresponds to the seven canonical networks. **C**. Lateral and medial views of the ALANs networks colored according to Yan and colleagues’ seven-network parcellation. For VISU, 100% of the regions pertained to the canonical visual network (violet). For SomatoMotor, 72% (blue) of the regions were in the Som/Motor network. For the ParietoFrontal, 80% of the regions corresponded to the Control network. By contrast, for PosteriorMedial and TemporoFrontal networks, the distribution across the seven canonical networks is more scattered. View of 3D white surfaces rendering on the BIL&GIN display template in the MNI space.

### The ParietoFrontal Network: A Central Role in Goal-Directed Orientation and Executive Control of Attention

Our findings demonstrate a significant overlap of the ParietoFrontal network with the Control network (Figure 4), particularly in regions encompassing the dorsolateral and superior medial prefrontal cortex and the inferior parietal cortex. This overlap underscores the ParietoFrontal network’s integral role in a variety of executive functions, including cognitive control, attention regulation, and working memory, resonating with descriptions of similar frontoparietal networks in the literature (Uddin et al., 2019; Vincent et al., 2008). The observed lateralization in the ParietoFrontal network aligns with studies suggesting hemisphere-specific roles: the right hemisphere’s involvement in attentional control and inhibition (Aron et al., 2004; Spagna et al., 2020) and the left hemisphere’s dominance in abstraction and hierarchical control, probably associated with the language processes (Nee, 2021). The lateralization of the parietofrontal network in visuospatial processes highlights its role in interhemispheric balance, being right-lateralized when interacting with attentional regions and left-lateralized with language regions, as shown by Wang and colleagues (Wang et al., 2014). Moreover, the intersection of the ParietoFrontal network with the dorsal attention network, particularly in right-hemispheric regions like the precentral sulcus and intraparietal sulcus, further highlights the ParietoFrontal network’s involvement in attentional orienting, consistent with the task demands of the line bisection judgment task. This finding is bolstered by graph theory analysis, identifying the inferior frontal sulcus (f2_2) as a hub node, likely mediating between the dorsal orienting and frontoparietal control systems. The role of the right middle frontal cortex (f2_2 and prec_1) as a link between ventral and dorsal networks, as suggested by resting-state functional connectivity studies (Fox et al., 2006), further supports this integrated perspective on attentional control. These results collectively reinforce the concept of lateralized control processes during visuomotor tasks, illuminating the complex interplay of cognitive control and attentional orienting networks in the brain.

### Interhemispheric Integration and Attentional Roles of the TemporoFrontal Network

In our study, the TemporoFrontal network stands out for its unique composition, encompassing both rightward temporal-frontal regions and leftward superior temporal cortex and subcortical nuclei. This bi-hemispheric characteristic positions it as a distinctly interhemispheric network. This alignment with studies on resting-state activity, which often group both left and right temporal regions, underscores the network’s involvement in detecting and reorienting attention toward salient stimuli (Menon & Uddin, 2010; Seeley et al., 2007).

Specifically, the inclusion of the left superior temporal gyrus region (T1_4), which exhibits language-related leftward asymmetry (Labache et al., 2019), suggests a broader functional scope for this network than previously recognized. Moreover, the detection of subcortical structures, particularly the thalamus, aligns with recent neuroanatomical models of the ventral (VAN) and dorsal (DAN) Attentional Networks, which emphasize the role of the pulvinar as a central region modulating information flow processing in attentional processes (Alves et al., 2022).

The right posterior temporal regions identified in our study are parts of the occipitotemporoparietal junction, contributing to a variety of behaviors and functions such as redirecting attention towards task-relevant stimuli within the VAN, self-perception, and social cognition (Corbetta & Shulman, 2002; Saxe & Kanwisher, 2003). Similarly, the right inferior frontal cortex is implicated in diverse cognitive functions, including the inhibition component of the VAN and social cognition. Numerous studies have aimed to delineate the functional subdivisions of these regions using task-based or large-scale network mapping approaches (Geng & Vossel, 2013; Igelström & Graziano, 2017). For example, recent research by Numssen and colleagues proposed an anterior/posterior functional specialization of the inferior parietal lobe across attentional, semantic, and social cognitive functions, as well as hemispheres (Numssen et al., 2021). Additionally, a coactivation-based parcellation of the right inferior frontal gyrus (IFG) revealed a complex functional organization. This organization includes a posterior-to-anterior axis, with action/motor-related functions concentrated in the posterior region and cognition/abstract-related functions in the anterior region. Moreover, a dorsal-to-ventral axis within the posterior IFG corresponds to distinctions between action execution and inhibition, while a similar axis within the anterior IFG delineates reasoning and social cognition functions (Hartwigsen et al., 2019). The rightward regions clustered in the TemporoFrontal network likely underlie the bottom-up attentional processes and inhibition required to perform the LBJ task.

Moreover, this complexity is also reflected in the overlap with the 7-networks parcellation (Figure 4), with 44% of the TemporoFrontal network overlapping with the default-mode network (DMN) and 33% with the VAN/Sal network. While the VAN is implicated in reorienting attention to salient stimuli in the environment, particularly when they are unexpected or novel, the Salience network (SN or Sal) is primarily involved in detecting and filtering salient stimuli from the environment that are biologically or emotionally relevant and require immediate attention (Seeley et al., 2007). The SN plays a key role in switching between different brain networks, facilitating the transition from the DMN to the frontoparietal executive network in response to salient stimuli, and includes regions such as the anterior cingulate cortex (ACC), the anterior insula, and parts of the dorsomedial prefrontal cortex (dmPFC). Those two networks share common brain regions, especially the ventral anterior insula. The anterior insula has been also shown to be a key region of the cingulo-opercular network (CON, (Dosenbach et al., 2006)). The CON is involved in maintaining task sets, sustaining attention, and cognitive control processes. It includes regions such as the anterior insula, the dorsal anterior cingulate cortex (dACC), the anterior prefrontal cortex, and the operculum. The CON is engaged in tasks requiring sustained attention, response inhibition, and error monitoring. It is associated with maintaining stable cognitive states and regulating attentional processes over time. Our analysis of the TemporoFrontal network’s asymmetry during the visuospatial task supports its involvement in these complex attentional mechanisms. Notably, the network’s hubs in regions like the anterior insula suggest a potential interaction site between the VAN, CON and SAL networks and also with the DAN (Cazzoli et al., 2021), underscoring its critical role in modulating attentional processes. Finally, the strong positive correlation observed between the ParietoFrontal and TemporoFrontal networks further emphasizes their collaborative function in attentional control, although further research is needed to fully elucidate the lateralization and functional dynamics of these high-order networks.

### Functional Integration and Spatial Processing in the PosteriorMedial Network

As identified in our study, the PosteriorMedial network encompasses a range of regions in the right hemisphere, including the posteromedial wall from the precuneus through the medial temporal lobe to the anterior temporal pole. These regions predominantly involve spatial cognition, attention, and memory (Cavanna & Trimble, 2006; Richter et al., 2019; Shulman et al., 2010). Notably, the right precuneus and posterior parietal cortex have been shown to exhibit a rightward bias during visuospatial tasks (Mahayana et al., 2014), suggesting their significant involvement in spatial processing. Compared to the 7-networks parcellation from Yeo and colleagues (Yeo et al., 2011), the regions within the PosteriorMedial network show a diverse overlap across multiple resting-state networks, including visual, dorsal attention, control, and default mode networks. This complex overlap pattern resonates with recent findings that identified intricate hippocampal-parietal circuits and connections to the parietal memory network (Seoane et al., 2022; Zheng et al., 2020), further supporting the involvement of the PosteriorMedial network in goal-oriented processing and stimulus recognition. Moreover, connectivity studies, such as those by Zhang and Li (Zhang & Li, 2012), demonstrate that the dorsal precuneus within this network exhibits strong connections with occipital and posterior parietal cortices and areas related to motor execution and visual imagery. This rich connectivity underscores the network’s role in integrating spatial, motor, and visual information. Regarding network correlations, the PosteriorMedial network showed the lowest overall connectivity, with a positive correlation with the VISU network and a slight negative correlation with the TemporoFrontal Network, highlighting its distinct functional profile. These findings emphasize the unique positioning of the PosteriorMedial network in the neural architecture, playing a pivotal role in spatial processing and integrating diverse cognitive functions.

### The Local Visual and SomatoMotor Networks

In our exploration of local visual (VISU) and sensorimotor (SomatoMotor) networks during the line bisection judgment task, we observed distinct patterns of BOLD asymmetry that align with existing literature on visuospatial attention and sensorimotor processing. Specifically, the VISU network demonstrated a pronounced rightward BOLD lateralization, independent of stimulus asymmetry, reflecting the engagement of top-down attentional processes and lateralized modulation of visual cortical regions, consistent with the interactions between the dorsal attention system and the visual occipital cortex (Corbetta & Shulman, 2002; Meehan et al., 2017).

Furthermore, our analysis reveals a robust collaboration between the SomatoMotor and VISU networks, as evidenced by their strong positive temporal correlation in mean intrinsic connectivity. This finding underscores the integrated function of these networks in visuomotor coordination, supporting the hypothesis of their cooperative role in complex cognitive tasks (Rizzolatti & Matelli, 2003). Additionally, we identified leftward areas overlapping with the dorsal attention network in the left hemisphere (Corbetta & Shulman, 2002; Petit et al., 2009), suggesting a significant role of the left hemisphere in coordinating eye movements in right-handed individuals. This observation, coupled with our findings of leftward asymmetries in regions associated with hand and mouth movements, illustrates the multifaceted nature of the left hemisphere’s involvement in visuospatial attention and motor planning in right-handers, underlining its intricate role in integrating eye, hand, and mouth movements during cognitive tasks.

### Evaluating ALANs Within the Broader Spectrum of Brain Lateralization Function

We here compared ALANs to a set of three other atlases we previously developed (Figure 5). These atlases have all been developed using the same methodology as in the present paper, with all having the purpose of characterizing the anatomo-functional support of lateralized cognitive brain function.

**Figure 5.**
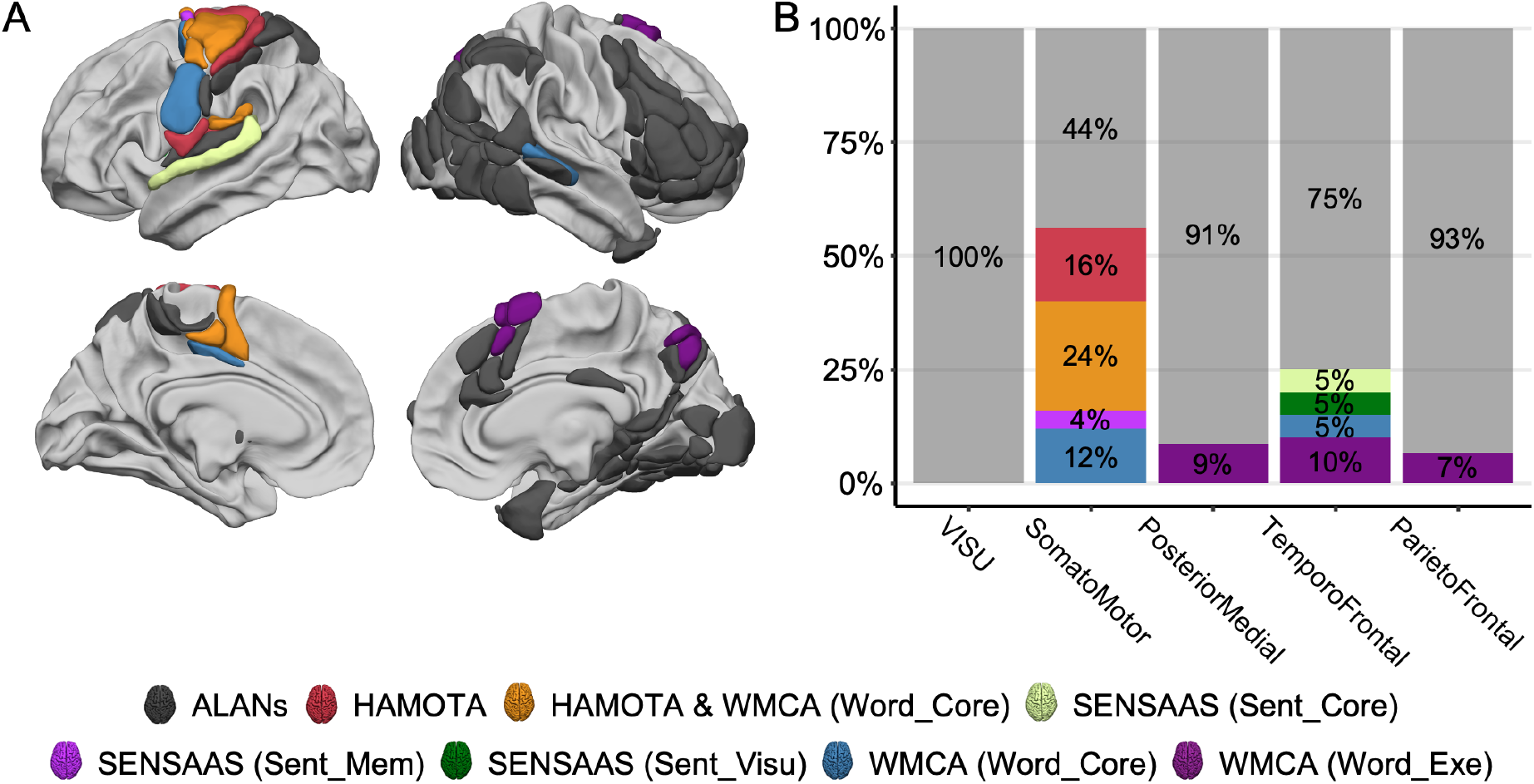
Comparison between the five ALANs (Atlas of Lateralized visuospatial Attentional Networks) clustered networks and three other functional atlases; HAMOTA: HAnd MOtor Area atlas (Tzourio-Mazoyer et al., 2021), SENSAAS: Sentence Supramodal Areas Atlas (Labache et al., 2019), WMCA: Word-list Multimodal Cortical Atlas (Hesling et al., 2019). **A**. Right and left lateral and medial views of the ALANs atlas. Regions are colored according to the HAMOTA, SENSAAS, and WMCA parcellations. View of 3D white surfaces rendering on the BIL&GIN display template in the MNI space. **B**. Repartition of the regions of the ALANs five-network parcellation across HAMOTA, SENSAAS, and WMCA.

Unlike the VISU network, which is exclusively linked to visual processes in the line bisection judgment task, the SomatoMotor network shows broader cognitive involvement. Specifically, 56% of the SomatoMotor network was found to be non-specific to visuospatial attention, suggesting its engagement in a wider range of cognitive functions. Specifically, the SomatoMotor network demonstrated significant overlap with the HAMOTA (HAnd MOtor Area atlas, (Tzourio-Mazoyer et al., 2021)), WMCA (Word-list Multimodal Cortical Atlas, (Hesling et al., 2019)), and SENSAAS (Sentence Supramodal Areas Atlas (Labache et al., 2019)) atlases (Figure 5). This overlap indicates a strong leftward asymmetry in regions associated with somatomotor response production. This asymmetry extends from primary and secondary somatosensory cortices to motor areas (Figure 5), highlighting the left hemisphere’s dominant role in processing and executing right-hand response production (Tzourio-Mazoyer et al., 2021) and coordinating subvocal articulation associated with finger selection (Hesling et al., 2019). Subvocal articulation particularly takes place in the Rolandic fissure (rol1), the only region overlapped by WMCA (Figure 5, (Hesling et al., 2019)) and involved in the mouth, larynx, tongue, jaw, and lip movement. The precuneus region of the PosteriorMedial network only overlaps with the executive network of WMCA (Figure 5), highlighting its role in mental imagery and/or episodic memory encoding related to the line bisection judgment task (Cavanna & Trimble, 2006). Concerning the ParietoFrontal network, the supplementary frontal eye field (SMA1, Figure 5) is also related to the executive network of WMCA, highlighting its role in evaluating value-based decisions involved in the line bisection judgment task (So & Stuphorn, 2012). Finally, the TemporoFrontal network had 25% of its regions overlapping with either SENSAAS or WMCA (Figure 5). Among them, two supplementary motor areas (SMA2 and SMA3) were related to the executive network of WMCA and the supplementary frontal eye field. The superior temporal sulcus (STS3), also known as the posterior human voice area (Pernet et al., 2015), was also found to be a key region in the core network of WMCA. This region is a key area in the interhemispheric communication processes, intertwining between prosodic and phonemic information (Hesling et al., 2019). Two regions overlapped with SENSAAS: the putamen (PUT_3), supporting executive functions and task monitoring in the processing of multimodal language processing (Labache et al., 2019; Monchi et al., 2006), and the superior temporal gyrus (T1_4) supporting amodal semantic combinations (Labache et al., 2019; Price, 2010).

As demonstrated in Figure 6, our analysis revealed a significant overlap (50%) between regions within the TemporoFrontal network and the homotopic version of the core multimodal sentence network (Labache et al., 2019). Notably, the pars triangularis of the inferior frontal gyrus (F3t) and the superior temporal sulcus (STS4), both key hub regions for the TemporoFrontal network, were also hubs for the core network of SENSAAS (Labache et al., 2019). This suggests a mirror-like organizational similarity between visuospatial attention and language processing networks, with the peripheral regions of the hubs probably defining the type of processes each hemisphere carries out. Similarly, the inferior frontal sulcus (f2_2), a hub for the ParietoFrontal network, was also a central region in the core network of SENSAAS and is on the verge of being a hub (Labache et al., 2019). Furthermore, recent findings reveal that with aging, language processing regions in the left hemisphere shift from leftward asymmetry to bilateral organization, similarly affecting the mnemonic regions’ symmetry (Roger et al., 2023). This reorganization suggests a nuanced interplay between language, memory, and visuospatial attention over time.

**Figure 6.**
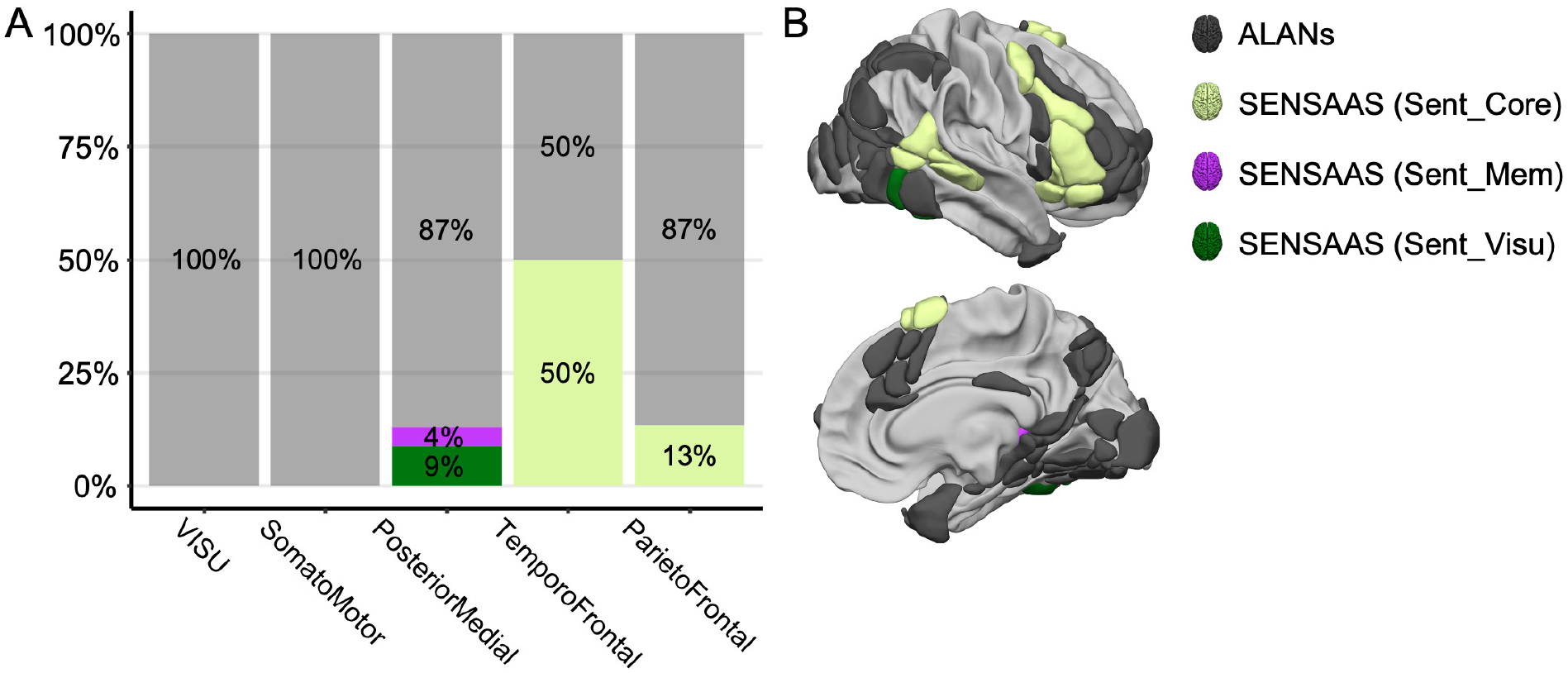
Comparison between the 5 ALANs (Atlas of Lateralized visuospatial Attentional Networks) clustered networks and the homotopic version of SENSAAS; Sentence Supramodal Areas Atlas (Labache et al., 2019). **A**. Repartition of the regions of the ALANs five-network parcellation across the homotopic version of the SENSAAS atlas. **B**. Right lateral and medial views of the ALANs atlas. Regions are colored according to the homotopic version of the SENSAAS parcellations. View of 3D white surfaces rendering on the BIL&GIN display template in the MNI space.

The other atlases of lateralized brain functions mentioned in this section are available to the community here: https://github.com/loiclabache.

## Conclusion

Our study elucidates the lateralized brain networks involved in visuospatial attention among right-handed individuals, highlighting the critical roles of the ParietoFrontal and TemporoFrontal networks. The discovery of significant overlaps with the contralateral sentence network emphasizes a complex interplay between attentional and language processes, shedding light on the brain’s functional asymmetry. These insights advance our understanding of cognitive function lateralization and pave the way for future research into atypical brain organization and hemispheric complementarity, with broad implications for both neuroscience and clinical practice. The homotopic Atlas of Lateralized visuospatial Attentional Networks (ALANs) is publicly available as a resource for future studies (Labache, 2024) and can be found here: https://github.com/loiclabache/ALANs_brainAtlas.

## Data and Code Availability Statement

The data, the code, and the atlas used to produce the results can be found here (Labache, 2024): https://github.com/loiclabache/ALANs_brainAtlas.

## CRediT Authorship Contribution Statement

**Loïc Labache:** Conceptualization, Data Curation, Formal Analysis, Investigation, Methodology, Software, Validation, Visualization, Writing - original draft, Writing - review & editing, Supervision, Project administration. **Laurent Petit:** Investigation, Resources, Writing - review & editing, Funding acquisition. **Marc Joliot:** Investigation, Resources, Writing - review & editing, Funding acquisition. **Laure Zago:** Conceptualization, Validation, Investigation, Resources, Visualization, Writing - original draft, Writing - review & editing, Supervision, Project administration, Funding acquisition.

## Acknowledgments

We deeply thank Dr. Nathalie Tzourio-Mazoyer for her thoughtful help in discussing the results. The results are part of the BIL&GIN database (Mazoyer et al., 2016).

